# *De novo* mutation rates in sticklebacks

**DOI:** 10.1101/2023.03.16.532904

**Authors:** Chaowei Zhang, Kerry Reid, Arthur F. Sands, Antoine Fraimout, Mikkel Heide Schierup, Juha Merilä

**Affiliations:** Area of Ecology & Biodiversity, School of Biological Sciences, The University of Hong Kong, Hong Kong SAR; Bioinformatics Research Centre, Aarhus University, Denmark; Research Program in Organismal & Evolutionary Biology, Faculty Biological and Environmental Sciences, University of Helsinki, Finland

**Keywords:** mutation rate, divergence time, genetic diversity, germline mutation rate, ninespine stickleback

## Abstract

Mutation rate is a fundamental parameter in population genetics. Apart from being an important scaling parameter for demographic and phylogenetic inference, it allows one to understand at what rate new genetic diversity is generated and what is the expected level of genetic diversity in a population at equilibrium. However, except for well-established model organisms, accurate estimates of *de novo* mutation rates are available for a very limited number of organisms from the wild. We estimated mutation rates (*µ*) in two marine populations of the nine-spined stickleback (*Pungitius pungitius*) with the aid of several 2- and 3-generational family pedigrees, deep (>50×) whole genome re-sequencing and a high-quality reference genome. After stringent filtering, we discovered 295 germline mutations from 106 offspring translating to *µ* = 4.64 × 10^−9^ and *µ* = 4.08 × 10^−9^ per base, per generation, in the two populations, respectively. Twenty percent of the mutations were shared by full-sibs showing that the level of parental mosaicism was relatively high. Since the estimated *µ* was 3.2 times smaller than the commonly used substitution rate, recalibration with *µ* led to substantial increase in estimated divergence times between different stickleback species. Our estimates of *de novo* mutation rate should provide a useful resource for research focused on fish population genetics and that of sticklebacks in particular.

## Introduction

Although much of the short-term evolution and adaptation is likely based on standing genetic variation (Barrett and Schluter 2008), new mutations are the ultimate source of genetic diversity. The rate at which new mutations arise is a key parameter in evolutionary biology and population genetics (Hartl and Clark 2006), but at the same time difficult to quantify as per-generation mutation rates are low (Lynch 2010). Traditionally, mutation rates (*µ*) have been estimated with the aid of locus-specific rates on the basis of phenotypes observed in crosses and pedigrees (e.g. Stadler 1930), mutation accumulation experiments (Mukai 1964) or inferred from sequence divergence among taxa (Kimura 1968). All of these approaches make assumptions that are known to be frequently violated, and consequently, they can provide only gross approximations of *de novo* mutation (DNM) rates (Smeds et al. 2016).

The drop in DNA-sequencing costs combined with improved variant calling methods have led to replacement of traditional approaches for mutation rate estimation with direct estimates obtained from DNA-sequence data (supplementary table 1 and fig. 1). However, direct estimation of DNM rates is not easy. Mutations occur at slow rates and each DNM has only 50% probability to be transmitted from a parent to offspring, and as such, a relatively large number of individuals from sequential generations need to be sequenced to have high detection probability. Even if enough DNMs can be confidently called, converting these to per generation (and year) mutation rates requires that the callable part of the genome (denominator of the rate estimate) is well estimated, which in turn requires a high-quality reference genome assembly (Besenbacher et al. 2015, 2019; Bergeron et al. 2022). In addition, to distinguish true DNMs from somatic mutations, controlling for false positive DNMs calls, each individual needs to be sequenced to high depth of coverage (Besenbacher et al. 2015; Bergeron et al. 2022). This means that mutation rate estimation is still costly for organisms with large genomes and not feasible for organisms lacking good quality reference genomes, against which sequenced reads can be confidently mapped. Furthermore, the mappable fraction of the genome should not be too small and it should be well defined (Bergeron et al. 2022). Therefore, direct estimates of mutation rates are mostly available from model organisms with well-developed genomic resources and typically from unnatural captive or laboratory colonies (e.g. Keightley et al. 2014; Milholland et al. 2017; Lindsay et al. 2019; Wang et al. 2020; Bergeron et al. 2021). Recently, estimates have started to become available for a limited number of non-model organisms such as cats, wolves, birds and the duck-billed platypus (e.g. Keightley et al. 2015; Smeds et al. 2016; Martin et al. 2018; Koch et al. 2019; Yang et al. 2021). However, the pedigrees in most of these studies have been small, typically comprising a dozen of individuals or less (e.g. Smeds et al. 2016; Besenbacher et al. 2019; Koch et al. 2019).

**Fig. 1.**
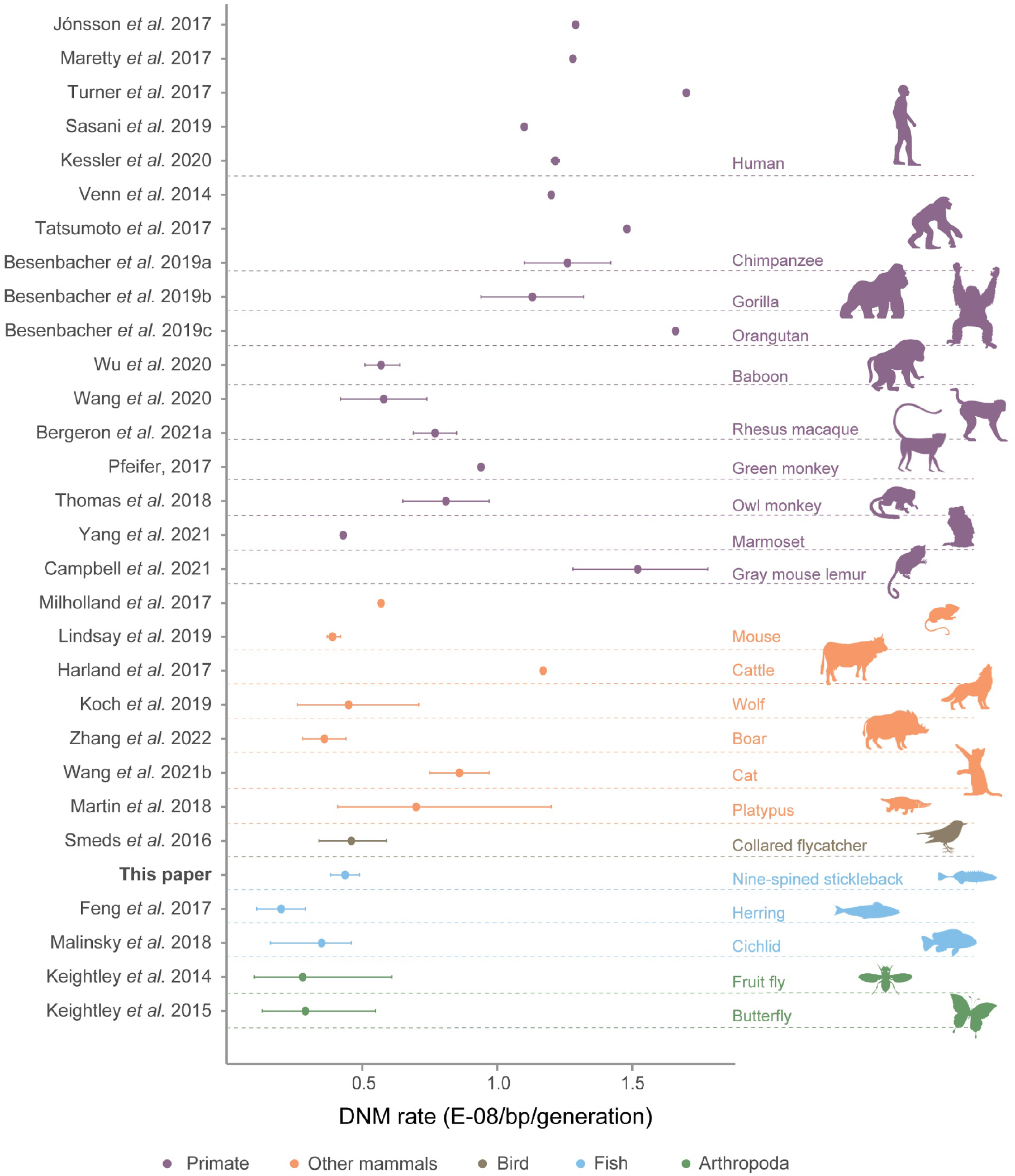
DNM rates to date. Per-site-per-generation DNM rate (10^−8^) estimates from studies which have used pedigree-based mutation rate estimation. Rates with no error bars do not have 95% confidence intervals available. For more details, please see supplementary table 1.

The nine-spined stickleback (*Pungitius pungitius*) is a small teleost fish which has recently been subject to many population genomic, demographic and phylogenetic investigations (e.g. Guo et al. 2019; Natri et al. 2019; Varandhajan et al. 2019; Yamasaki et al. 2020; Fang et al. 2021; Kemppainen et al. 2021; Wang Y et al. 2022; Feng et al. 2022; Kivikoski et al. 2023), meaning that there is a community of researchers that would benefit from access to DNM rates in this species. This is because mutation rate estimates are key scaling parameters in many population genetic, demographic and phylogenetic inferences (e.g. Koch et al. 2019). Therefore, many-fold differences, for instance in estimates of migration rates, effective population sizes (*Ne*), genetic diversity and divergence times among taxa, can ensue if inaccurate estimates of *µ* are used to derive them (Besenbacher et al. 2019; Koch et al. 2019; Tiley et al. 2020). In fact, studies of sticklebacks have so far resorted to using the substitution rates between three- (*Gasterosteus aculeatus*) and nine-spined stickleback (Guo et al. 2013) and an estimate of divergence time as proxy of *µ* (e.g. Liu et al. 2018; Ravinet et al. 2018; Varadharjan et al. 2019; Yamasaki et al. 2020; Dahms et al. 2022; Feng et al. 2022).

Here, we aimed to obtain accurate estimates of DNM rates for outbred nine-spined sticklebacks using a high-quality reference genome assembly (Kivikoski et al. 2021) and deep (50×) sequencing of multigenerational pedigrees (2- and 3-generations) consisting of a total of 128 individuals from two marine populations (five families from each) separated by distance of over 300 km. In addition, we investigated where in the genome the DNMs occurred and whether they were associated with specific genomic features. Finally, we utilised these estimates to assess the consequences of using mutation rates instead of fossil calibration points and substitution rates to estimate divergence times among different stickleback species and lineages.

## Material and Methods

### Sampling

Twenty-two sexually mature male and female nine-spined sticklebacks forming the F_0_ generation were sampled with beach seine nets from May to June 2018 from Pori (POR; 61.591°N, 21.473°E) and Tvärminne (TVA; 59.833°N, 23.200°E) in Finland. Both localities are Baltic Sea coastal sites, and hence, the parental generation originated from outbred marine populations. The fish were transported to the aquaculture facility of Viikki Campus (University of Helsinki) and maintained in 17°C in aerated aquaria until used in artificial fertilisations (for details of rearing conditions and procedures see Fraimout et al. (2022). For each of the two marine populations, five 2- or 3-generational pedigrees were produced from the wild-caught F_0_ individuals by artificial crossing, where the last generation of each pedigree consisted of 10 full-sibs (fig. 2). Briefly, *in vitro* crosses were performed by squeezing eggs from females and combining them with minced testes dissected from males (euthanised with MS-222) in a Petri dish, where gametes were mixed gently to ensure fertilisation. The resulting clutches were first reared in Petri dishes until they hatched and the fry started independent feeding. The F_1_ generation fish were reared for approximately 400 days (mean: 403.6 days), after which they were euthanised (using MS-222) or kept for breeding the F_2_ generation. In the case of the families where an F_2_ generation was produced, artificial crosses of F_1_ parents were performed as described above to produce F_2_ offspring, the latter which were euthanised (with MS-222) and preserved in ethanol two days post-hatching. All euthanised individuals (across generations) were stored in 95% ethanol to preserve DNA for extractions. Altogether, this study included 128 individual fish; 12 POR and 10 TVA F_0_ wild caught specimens, 34 POR and 42 TVA F_1_ specimens and 20 POR and 10 TVA F_2_ specimens (Total n_POR_ = 66, n_TVA_ = 62). Hence, the number of trios (i.e. groups of two parents and their offspring) for POR and TVA were 54 and 52, respectively.

**Fig. 2.**
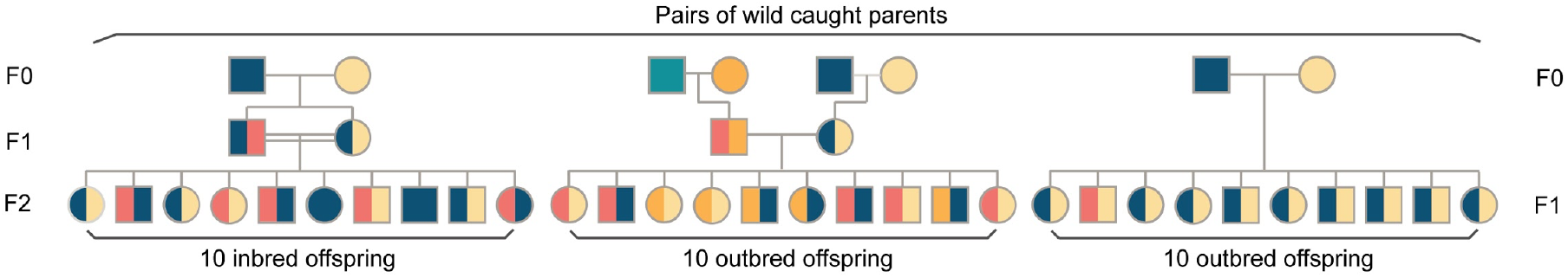
Pedigree types. Three types of pedigree structures used in this study: three-generation inbred line (left), three-generation outbred line (middle) and two-generation outbred line (right). Squares and circles represented males and females respectively. The mutated haplotypes are shown in red and the normal wild type haplotypes in other colours.

### DNA extraction and Sequencing

Genomic DNA was extracted from fin clips using a modified salting-out protocol as described by Sunnucks and Hales (1996). The DNA purity of samples was evaluated with a NanoDrop spectrophotometer and the concentrations were quantified using Qubit dsDNA HS Assay Kit with Qubit™ 4.0 (Invitrogen, CA, USA).

For TVA samples, the genomic DNA libraries were constructed with the NEBNext® Ultra™ II FS DNA Library Prep Kit (Agilent, CA, US). Thereafter, qPCR quantification of the sequencing libraries was conducted following the NovaSeq v1.5 protocol at the Biomedicum Functional Genomics Unit (FuGU) of the Helsinki Institute of Life Science (HiLIFE) and Biocenter Finland (BF) research infrastructure. DNA samples (1 *µ*g) for each individual of the POR population were sent to Beijing Genomics Institute (BGI) for PCR-free library construction using their proprietary DNBseq platform for NGS sequencing. All samples were whole-genome-sequenced to 50× target coverage.

### Read mapping and variant calling

The paired-end data was processed following Feng et al. (2022). In brief, the raw reads were mapped to the most recent available nine-spined stickleback reference genome (version 7, GCA_902500615.3, Kivikoski et al. 2021) using the Burrows-Wheeler Aligner (BWA) with mem option (v0.7.17; Li 2013). Aligned reads were then sorted and indexed with mate coordinates flagged through SAMtools v1.10 (Li et al. 2009). The duplicate reads were marked with PicardTools (v2.18; http://picard.sourceforge.net). Base-quality score recalibration (BQSR) was also performed in GATK (v4.2.2.0; Van der Auwera et al. 2020) using hard filtered SNPs and indels.

Following the best practices workflow of GATK (Poplin et al. 2017), the nucleotide variants were called using HaplotypeCaller in ERC mode, with several annotations being added (e.g. “MappingQuality”, “FisherStrand”, etc.) for downstream filtrations. The per-sample VCF files were then jointly genotyped by the CombineGVCFs and GenotypeGVCFs modules for each population and for each parent-offspring trio.

### Pedigree examination

To confirm genetic relationships (e.g. parent-offspring) and structure of the pedigrees, several analyses were performed before germline mutation identification (where sex chromosome (LG12) and unassigned contigs were excluded). We firstly estimated the probabilities of identity-by-descent (IBD) for each pedigree with PLINK (v1.90; Chang et al. 2015). The Z_0_:Z_1_:Z_2_ (probabilities of sharing no, one and two alleles) for parent-offspring relationship should be close to 0:1:0, whereas that for full-sibs should be close to 0.25:0.5:0.25. We then also performed parentage analyses to check if the paired parents could be correctly assigned back to their offspring in FRANz (v1.9.999; Riester et al. 2009). The 012 matrices were generated in VCFtools (v0.1.16; Danecek et al. 2011) by allowing 10% missing genotype data and a minor allele frequency (MAF) of 0.01. Finally, a principal component analysis (PCA) for all individuals was performed to check that the individuals from the same pedigree would cluster together in a PCA plot. The PCA plots were generated with ANGSD and PCANGSD (v0.939; Korneliussen et al. 2014). These checks confirmed that genetic relationships among the sequenced individuals were as assumed.

### Identifying the candidate de novo mutations (DNM)

For each parent-offspring trio (n = 106), the variants in each trio VCF file were filtered to a subset of single nucleotide variants (SNVs) by BCFtools (v1.10; Danecek *et al*, 2021) based on the Mendelian violation (fig. 2): We considered an offspring heterozygous variant (0/1) to be a DNM when their parents were both homozygotes for either reference (0/0) or alternative allele (1/1). A series of site filters and individual filters were then applied to these SNVs following the “Mutationathon” guidelines (Bergeron *et al*, 2022) which included:

#### (i) Site filtering

Following GATK best practice pipeline (Poplin et al. 2017), hard filtering was applied to all individuals removing the low-quality positions with the following parameters: Quality by depth (QD) < 2.0, Mapping Quality (MQ) < 40.0, Fisher’s exact test on strand bias (FS) > 60.0, Standard odd ratio (SOR) > 3.0, Mapping quality rank sum test (MQRankSum) < -12.5 and Read position rank sum test (ReadPosRankSum) < -8.0.

#### (ii) Individual filtering

DNM candidates were filtered based on the following criteria to eliminate the false positives: 1) sequencing depth (DP ≤ 20 and DP ≥ 100) and genotyping quality (GQ ≤ 80) for both parents and their offspring were examined to exclude genotyping errors or read misalignments in regions of high complexity; 2) an allelic depth filter (AD1 > 0 for 0/0 or AD0 > 0 for 1/1) was applied for the two parents to ensure they are real homozygotes; 3) filters of allelic balance (AB < 0.3 and AB > 0.7) and sequencing depth (DP < 0.5DP_trio_ and DP > 2DP_trio_) were applied for offspring to confirm they are true heterozygotes; 4) a filter to remove clustered sites where more than one DNM candidate were observed within 100 bp as multiple, adjacent mutations are expected to occur in extremely low probabilities; 5) an inspection to delete DNM candidates within 5 bp away from any indels to avoid any uncertainties brought by the realignment step; 6) deletions of DNM candidates that occurred repeatedly in multiple unrelated samples, but keep a separate check of those shared among full-sibs.

### *De novo* mutation rate estimation

The per site per generation mutation rate (*µ*) for each offspring was calculated as:

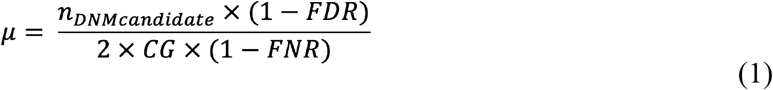

The number of callable genome sites (CG) was obtained by applying a sequencing depth (DP) filter on the read alignments (bam files) in a given trio. The sites in an offspring were counted as ‘callable’ only if they were with more than half and less than double of the total DPs within its trio family (0.5DP_trio_ < DP_child_ < 2DP_trio_).

Assuming the true heterozygotes (0/1) in each offspring were those where one parent carried 0/0 and another had 1/1, we considered the false negative rate (FNR) for each offspring as the percentage of true heterozygotes that were purged by the abovementioned AB filter (AB < 0.3 and AB > 0.7).

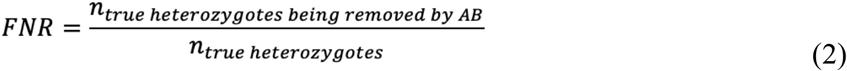

False positives were further identified manually using visualisation by IGV (Thorvaldsdóttir et al. 2013), where the genomes from each trio set were checked at the same time to ensure that the raw reads supported each genotype, and only those that were well-supported were retained. Finally, the candidate DNMs were removed if, firstly, both or one of the parents carried the same mutation as their offspring (supported by up to 10% or more raw reads in sum) or, secondly, the offspring was incorrectly identified as a heterozygote based on poor mapping in the positions around to candidate DNM. The false discovery rate (FDR) was then calculated from:

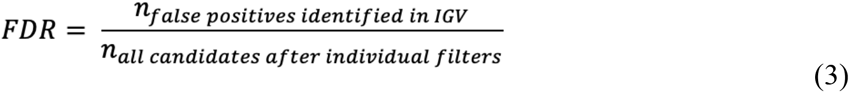

The mutation rates were estimated for the two populations separately, but as they were not significantly different (see Results), we also combined all families to single analysis to gain accuracy in our estimate of *µ*.

### Mutation spectrum and genomic context analyses

Although DNMs are usually distributed randomly throughout the genome, they typically show distinct frequencies in relation to which mutation spectrum they belong to (Milholland et al. 2017; Sasani et al. 2019; Wang et al. 2020). For example, mutation rates are observed to be particularly elevated in CpG sites where more deamination of methylated cytosines appears to occur (Razin and Riggs 1980; Zemojtel et al. 2011; Kong et al. 2012; Milholland et al. 2017; Wang RJ et al. 2022). First, mutation spectra were analysed based on alternative and reference alleles in vcf files. Secondly, DNMs were divided into transversions (Tv: A:T>C:G, A:T>T:A, C:G>A:T, and C:G>G:C) and transitions (Ts: A:T>G:C and C:G>T:A). Thirdly, CpG islands (CGIs) were predicted by applying the “twoBitToFa’’ program (http://genome.ucsc.edu/cgi-bin/hgTrackUi?g=cpgIslandExt, Miklem and Hillier; unpublished) to the reference genome following criteria outlined in Gardiner-Garden and Frommer (1987). The DNM rates of CGI and non-CGI areas were estimated separately for each offspring, where the callable genome sizes of both regions were calculated as described above. Finally, we annotated each mutation in relation to the genomic location (*viz*. within exon, intron or outside coding sequence) and mutation type (non-synonymous or synonymous) according to annotations in the previously published assembly of nine-spined stickleback (Version 6; Varadharajan *el al*. 2019) and a liftover file to Version 7 (Rastas 2020; https://sourceforge.net/p/lep-anchor/code/ci/master/tree/liftover.awk).

### Phylogenetic dating

To understand the impact of the estimated DNM rate on divergence time estimates, we reconstructed evolutionary relationships of Guo et al.’s (2019) dataset for RAD-seq data of stickleback lineages using the BEAST package (v.2.6.7; Bouckaert et al. 2019) with two different dating approaches. Herein, the input dataset consisted of 1,708 SNPs from 65 *Pungitius* individuals representing seven independent lineages, as well as of four *Gasterosteus* and two *Culaea* sticklebacks as outgroups. Specifically, we compared the divergence times of two phylogenies/evolutionary scenarios, selecting and setting different rate priors for dating: 1) Our estimate of the DNM rate – where we converted the per generation estimates of *µ* to per million year by assuming a generation length of two years (De Faveri et al. 2014); and 2) the synonymous substitution rate (SSR) between three- and nine-spined sticklebacks (7.1× 10^−9^/bp/yr; Guo et al. 2013) – which has been widely applied in literature to date (see Introduction). For congruence in each scenario, input files were constructed in BEAUti (BEAST package) using an optimised relaxed clock approach and the Yule tree prior. Therein to limit error and obtain the most accurate phylogeny, four independent runs of 100 million generations (sampling every 10,000 generations) were conducted as implemented in BEAST. They were then combined using LogCombiner (BEAST package) with 10% burnin (as assessed by parameter convergence in Tracer v.1.7.2; Rambaut et al. 2018). Tracer was again used to ensure that combined log files’ effective sample size (ESS) values were >200 in each scenario. TreeAnnotator (BEAST package) was then used for each scenario independently to summarise trees with no further burn-in for nodal support and date comparisons (supplementary fig. 4). Finally, the focal phylogenies presented were plotted with DensiTree which visualises the quantitative patterns across all trees (Bouckaert 2010).

## Results

### De novo mutation rates in nine-spined sticklebacks

A total of 1.17 million autosomal (and pseudoautosomal) variants passed the Mendelian violation. Herein, 534 putative DNMs were detected in POR and TVA families excluding those shared among siblings. After visualisation with the IGV tool, the number of DNMs were reduced to 160 and 135 for POR and TVA, respectively. The DNMs were widely dispersed throughout the genome (supplementary fig. 1). Power of all individual filters are reported in supplementary fig. 2 and no detectable batch effects were observed.

Based on read depths among trio families, the mean callable genome size was estimated to be 367.80Mb, 86.78% of the entire genome without sex chromosomes (but including the pseudoautosomal region, >16.9 Mbp on LG12) and unassigned contigs. The average false negative rate (FNR) was 5.81%. We inspected both the original and realigned (by choosing “-bamout” function) bam files for the variant calling step in IGV and found that the realignment procedure often led to disappearance and appearance of candidate DNMs. Thus, the manually curated FDR was 53.9% before and 21.9% after realignment in the GATK HaplotypeCaller. Following Bergeron et al. (2021), we eventually adopted the more conservative approach (the former one) which detected an average of 3.03 DNMs per individual (fig. 3a). Combining all the statistics above, the final estimate of single nucleotide germline mutation rate was 4.37 × 10^−9^/bp/generation (95% confidence interval (CI): 3.83–4.90 × 10^−9^). This translates to a yearly DNM rate of 2.18 × 10^−9^/bp/yr (CI: 1.92–2.45 × 10^−9^/bp/yr). There was no significant difference in DNM rate between the two populations (POR: 4.64 × 10^−9^, CI: 3.89–5.40 × 10^−9^ and TVA: 4.08 × 10^−9^, CI: 3.31–4.86 × 10^−9^; t-test: *t*_103.78_ = 1.03, p = 0.30; fig. 3b), the two sexes, pedigree types (inbred vs. outbred, fig. 2), or offspring generations (supplementary fig. 3a-c). The rates of DNMs transmitted from F_1_ to F_2_ were not statistically different between two pedigree types either (supplementary fig. 3d).

**Fig. 3.**
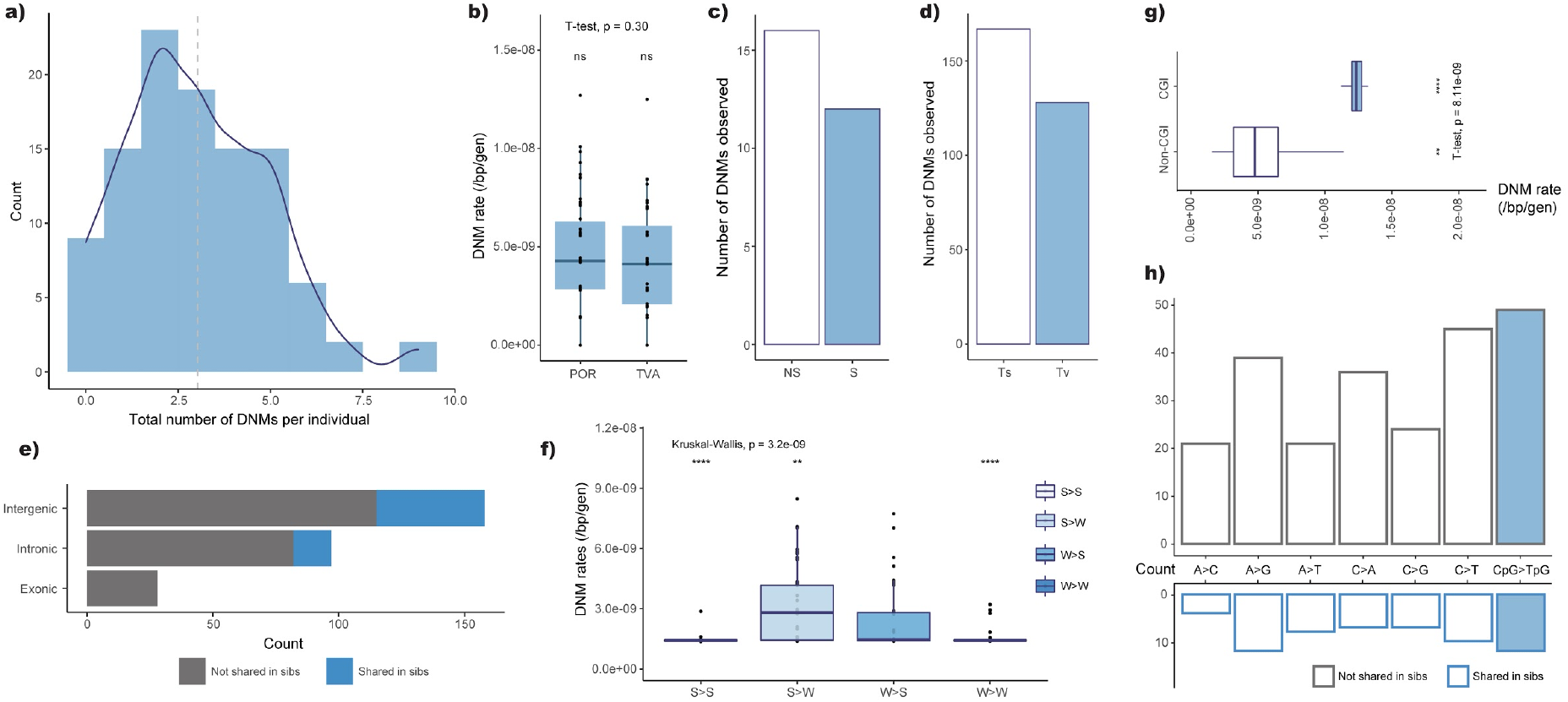
Mutation rates, types and spectra. a) Frequency distribution of DNMs detected per individual (dotted line is the mean = 3.03). b) Mean DNM rates in POR (4.64 × 10^−9^ [95% CI: 3.89 - 5.40 × 10^−9^]) and TVA (4.08 × 10^−9^ [95% CI: 3.31 - 4.86 × 10^−9^]) populations (t_103.78_ = 1.03, p = 0.30). c) Observed number of non-synonymous (NS) and synonymous (S) DNMs. d) Observed number of transversion (Tv) versus transition (Ts) mutations. e) Number of DNMs located in intergenic, intronic or exonic areas, categorised according to mutations being shared among full-sibs (blue) or not (grey). f) DNM rate of strong-to-weak-pairing type (S>W) mutations - note this was significantly higher (Kruskal-Wallis test) than in the other types (S>S: C>G, S>W: C>A or C>T, W>S: A>C or A>G, W>W: A>T. The respective mean DNM rates after corrected by FNR were: 1.47, 2.88, 2.39, and 1.61 × 10^−9^/bp/generation). g) Comparison of DNM rates in CpG island (1.53 × 10^−8^/bp/generation) and non-CpG island regions (5.05 × 10^−9^/bp/generation, t_25.47_ = 8.41, p = 8.11e-09). h) Mutation spectrum of the detected DNMs separated according to if mutations were shared among siblings (below, blue border) or not (above, grey border).

### Characterisation of mutation spectra

Among all DNMs, 56.6% were transitions (Ts) while 43.4% were transversions (Tv), showing a Ts:Tv ratio of 1.30 (χ^2^=71.92, df = 1, p < 2.2e-16; fig. 3d). The most common mutation type was C:G to T:A transition (116 out of the total 295), of which 52.6% were CpG > TpG mutations. We also observed a higher proportion of strong-to-weak pairing DNMs (S>W, C:G > A:T or C:G > T:A, 53.9%) with an average rate of 2.88 × 10^−9^/bp/generation. This DNM rate is significantly higher than the other types of substitutions in pairwise Wilcoxon test (vs. S>S: p = 1.4e-07; S>W: p = 1.7e-02; W>W: p = 1.4e-05; fig. 3f). In addition, 29 point mutations were observed within CpG islands accounting for 9.8% of all DNMs detected. We estimated a three-fold higher DNM rate in CpG islands (1.53 × 10^−8^/bp/generation) than in non-CpG islands (5.05 × 10^−9^; *t*_25.47_ = 8.41, p = 8.11e-09, fig. 3g).

Of the DNMs residing on the annotated parts of the v6 genome assembly (Varadharajan el al. 2019), 158 were within intergenic areas and 97 within introns, whereas 28 resided within gene coding sequences (CDS) and 12 in untranslated regions (UTRs, fig. 3e). There was no significant difference between the observed DNM frequencies and their expectation given the genomic coverage of each category (*viz*. intergenic, intronic or exonic; χ^2^ = 6.36, df = 3, p = 0.095). A total of 16 non-synonymous (NS) and 12 synonymous (S) exonic mutations were detected, among which 13 were CpG to TpG mutations (8 and 5 for NS and S DNM respectively). Furthermore, only one exonic DNM was found at the splicing sites which shifted the translation frames, potentially causing a loss-of-function (LOF) to the CDS. Except for this, no other LOF DNMs were detected including stop-codon variants.

In our dataset where the last generation of all pedigrees consisted of 10 full-sibs, 60 mutations, accounting for 20.3% of the total 295 DNMs, were carried by two or more siblings of the same parents suggesting that they had occurred during early germ cell divisions (parental mosaicism; Zlotogora 1998). These mosaic mutations only occurred in intergenic and intronic regions, but did not occur on exons (fig. 3e). Also, we did not detect any significant differences in mutation spectrum between shared and non-shared DNMs (χ^2^ = 2.92, df = 6, p = 0.82), including the fraction of CpG > TpG DNMs (20.0% vs. 20.8%; fig. 3h) or CGI variants (χ^2^ = 3.23, df = 1, p = 0.07).

### Divergence time estimation with DNM rates

Phylogenies, following the two dating approaches, generated in BEAST contained many well-supported nodes (supplementary fig. 4). Although there was one instance of branch swapping between the phylogenies (Node D; supplementary fig. 4), it is important to note that this node lacked significant support in both scenarios (PP ≥ 0.95) and otherwise the phylogenetic relationships were conserved. The dates of three nodes that most studies have focused on (i.e. A = divergence of *P. pungitius* and *G. aculeatus*; B = the MRCA for all *P. pungitius* lineages; C = divergence of eastern and western European lineages of *P. pungitius*; Guo et al. 2019; Feng et al. 2022) were shared and supported in both topological scenarios (fig. 4 and supplementary fig. 4), though these show key differences in dates depending on the method applied: The date comparison of scenarios using our Guo et al. (2013)’s SSR versus our DNM rate showed the divergence time of node A shifted from 5.6 Mya (95% highest posterior density (HPD): 3.3–8.1 Mya) to 18.3 Mya (95% HPD: 11.1–26.7 Mya), node B shifted from 2.6 Mya (95% HPD: 1.4–4.0 Mya) to 8.4 Mya (95% HPD: 4.7–13.0 Mya) and node C shifted from 1.3 Mya (95% HPD: 0.7– 2.0 Mya) to 4.2 Mya (95% HPD: 2.4–6.6 Mya) – equating to on average 2.23 times older dates across the three nodes based on our DNM rate (fig. 4).

**Fig. 4.**
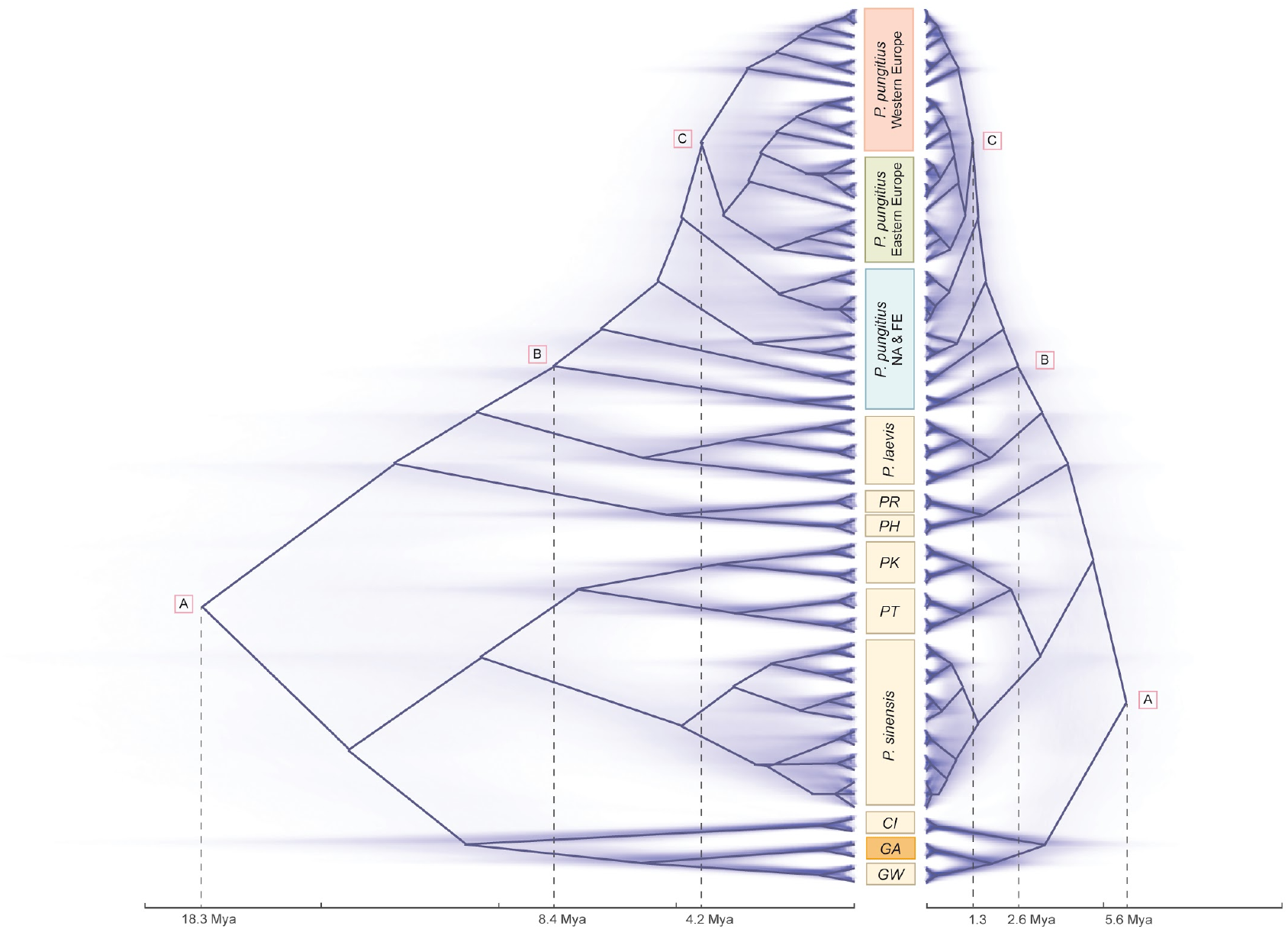
DNM and substitution rate-based phylogenies of *Pungitius* sticklebacks. DNM rate-based tree on left and substitution rate-based tree on right. Solid lines represent the summarised canal trees with maximum clade credibility scores, while the faint lines represent consensus trees for all topologies. (PR: *P. platygaster*, PH: *P. hellenicus*, PK:*P. kaibarae*, PT: *P. tymensis*, GW: *G. wheatlandi*, GA: *G. aculeatus*, CI: *Culaea inconstans*, FE: Far Eastern lineage, NA: North American lineage)

## Discussion

Although mutation rate is a fundamentally important quantity in evolutionary biology and genetics, accurate estimates for vertebrates beyond primates are still rare (supplementary table 1 and fig. 1). Here, we have provided a pedigree-based germline mutation rate estimate for sticklebacks based on, by far, the largest number of trios that any non-human study has used (supplementary table 1 and fig. 1). The estimated mutation rate for sticklebacks (0.437 × 10^−8^/bp/generation) is much lower (2.3–15.6×) than the rates that have been applied in earlier studies of sticklebacks (1.42 × 10^−8^, Guo et al. 2013; 6.8 × 10^−8^, Roesti et al. 2015; 3.7 × 10^−8^, Liu et al. 2016; and 1 × 10^−8^/bp/generation, Liu et al. 2018). Not surprisingly, application of the new mutation rate estimate established herein, as a prior when dating divergence times among stickleback clades, pushed the estimated divergences back in time quite considerably (fig. 4 and supplementary fig. 4).

To date, pedigree-based germline mutation rates have been estimated in 23 vertebrate species in 33 separate studies (supplementary table 1 and fig. 1). Most of these estimates come from studies of humans and primates (62.5%), followed by studies of other mammalian species (21.9%). Despite teleosts being the most species-diverse group of vertebrates (Venkatesh 2003), germline mutation rate estimates have only been estimated in four teleost fish species (supplementary table 1, shaded in blue). Compared to the estimates from the Atlantic herring (*Clupea harengus*, Feng et al. 2017) and three cichlid fish species (*Astatotilapia calliptera, Aulonocara stuartgranti*, and *Lethrinops lethrinus*, Malinsky et al. 2018), our estimate of germline mutation rate for nine-spined sticklebacks is 2.2 and 1.2 times higher, respectively, making the stickleback mutation rate the highest reported for teleost fishes, to date. In fact, our estimate is very similar to that of the only available estimate from birds *(Ficedula albicollis*) and is on par with those of grey wolf (*Canis lupus*), marmoset (*Callithrix jacchus*), baboon (*Papio anubis*) and rhesus macaque (*Macaca mulatta*; supplementary table 1 and fig. 1). However, with only four estimates of *µ* for teleost fishes based on small sample sizes (herring, four parents and 12 offspring, three cichlid species with one parent-offspring trio sequenced each) to date, any generalisations about mutation rates in fishes and their magnitude relative to other taxonomic groups requires additional pedigree-based estimates from different fish taxa to become available.

Low temperatures have been suggested to influence mutation rates due to slower metabolic rates (Martin and Palumbi 1993). Feng et al. (2017) discussed this as a possible factor explaining the low mutation rate in the Atlantic herring. While the higher mutation rates of Lake Malawi cichlids living in warmer waters align with this explanation (Malinsky et al. 2018), the even higher mutation rate estimates for nine-spined sticklebacks herein contradict it. Namely, the nine-spined sticklebacks used in our study originate from the Baltic Sea where sticklebacks are exposed to the same thermal conditions and metabolic constraints as Atlantic herring. Hence, the effect of environmental temperature on metabolic rates, and thereby to mutation rates, do not seem to be a likely explanation for low mutation rate in the Atlantic herring.

Parental age and gender are known to influence mutation rates in vertebrates. Our *µ* could be a slight underestimate compared to the situation in the wild if older parents generate and transmit more mutations to their offspring (e.g. Kong et al. 2012; Wong et al. 2016; Jónsson et al. 2017, 2018; Wang RJ et al. 2022). This is because our lab reared parents (F_1_ individuals in 3-generation families) were probably younger than their wild-caught parents. However, since more wild-caught (n=22) than F_1_ (n=6) parents were included into the analysis, any bias due to age variation is unlikely to be large. In fact, mutation rates estimated from wild-caught and F_1_ parents did not differ (*t*_71.92_ = 1.43, p = 0.16). Furthermore, one should also note that per year mutation rate estimates are subject to assumptions regarding the generation time used. For populations with overlapping generations, the generation time equals the mean age of parents (Hill 1979). Although we do not know the age of wild caught parents, the use of published estimates of the age of reproductive Baltic Sea sticklebacks (De Faveri et al. 2014) should provide a good proxy of the generation time for this species.

Fathers are known to generate and transmit more mutations to their offspring in primates (e.g. Kong et al. 2012; Wong et al. 2016; Jónsson et al. 2017, 2018; Wang et al. 2020; Wu et al. 2020), mice (Lindsay et al. 2019) and domestic cats (Wang RJ et al. 2022). Sperm is also generally more methylated than eggs, especially on CpG islands (Rahbari et al. 2016; Milholland et al. 2017), thus more DNMs is expected to be observed in chromatids inherited from fathers. Whether this applies also to fish has not been studied yet. In this study, we did not analyse whether there was paternal bias in *µ*, but since a similar number of parental males and females were analysed, this should not influence the overall *µ* estimate. An explicit test for sex-specific mutation rates in sticklebacks needs to await for a larger sample size of aged adults as used in this study.

One of the advantages of having direct estimates of germline mutation rates is that they allow one to probe long-term effective population sizes by substituting *µ* and nucleotide diversity (π) to solve effective population size (*N*_e_ = π/4*µ*, Watterson 1975). This gives an estimated long-term *N*_*e*_ for *P. pungitius* in the range of approximately 139,383–215,585 individuals. These estimates are an order of magnitude larger than estimates in Feng et al. (2022) obtained with coalescent methods (*N*_*e*_ ∼15,000–40,000). Yet, these numbers are likely to still be orders of magnitude lower than actual census population sizes of sticklebacks in the Baltic Sea. However, one has to remember that the *N*_*e*_ derived from the equation above refers to populations in mutation-drift equilibrium. In the case of Baltic Sea sticklebacks, the equilibrium assumption is likely to be violated due to post glacial population expansion and rampant introgression between divergent *P. pungitius* lineages (Feng et al. 2022). All these factors will influence π and thereby also the *N*_*e*_. In the same vein, the drift threshold *N*_*e*_’s obtained from the equation above would be overestimated if mutation rates over the last 1–2 Mya have been declining (Burridge et al. 2008).

Our analyses of mutation spectra in sticklebacks were largely congruent with those from mammalian studies (Pfeifer 2017; Koch et al. 2019). For example, we found over-representation of C>T transversions, more frequent weak-to-strong pairing mutations and random distribution of DNMs in intergenic, intronic and exonic areas. We also observed a high proportion of CpG>TpG mutations (20.68%), falling in the range observed in other species (9%–25%, Venn et al. 2014; Smeds et al. 2016; Thomas et al. 2018; Besenbacher et al. 2019; Campbell et al. 2021). Hence, these findings seem to suggest that the mutation process in fish bears close similarity to that in mammals. However, DNM rates in the CpG islands in our data exceeded that normally seen in mammals. Regions with high GC contents are supposed to have less CpG > TpG mutations and lower mutation rates than the other genomic regions (Youk et al. 2020). The gene PRDM9 specifies where recombination mediated double-stranded breaks occur in most mammals and likely also in other vertebrates (Cavassim et al. 2022). In canids and birds, in which PRDM9 has been lost, recombination is localised around “promoter-like” features and, in particular CpG islands, where mutations are more frequent. In three-spined sticklebacks, not all PRDM9 domains are present, and it is hypothesised that another mechanism determines recombination hotspots (Shanfelter et al. 2019). Therefore, a possible explanation for the high DNM rates detected in CGI regions of the genome of nine-spined sticklebacks could also be due to the lack of regulation from PRDM9.

We discovered that about 20% of DNMs were shared among full-sibs, suggesting that these mutations occurred post-zygotically at very early stages of development of the parental germline. This is a fairly high frequency compared to studies of humans (1.3%; Rahbari et al. 2016 and 3%; Sasani et al. 2019) and apes (3.5%; Bergeron et al. 2021), but a similar to a value estimated in mice (18%; Lindsay et al. 2019). However, a comparable estimate from the herring is much higher (50%), but this estimate is based on a very small sample size (Feng et al. 2017). Nevertheless, it appears as if parental mosaicism might be higher in fish than long-lived mammals.

Mutation rates are important in calibrating molecular clocks as well as in converting branch lengths of genealogies to units of time (Kimura 1968; Koch et al. 2019; Tiley et al. 2020). Hence, any uncertainty about mutation rates can directly propagate to distort demographic inferences, such as divergence times, effective population sizes and migration rates among populations (e.g. Ségurel et al. 2014; Koch et al. 2019). Our results provide a case in point: By calibrating the *Pungitius* phylogeny with our direct estimate of *µ* had a dramatic effect on divergence times pushing them back millions of years from the recent estimates (Fang et al. 2021; fig. 4). It is also worth noting that the divergence time estimates based on our *µ* aligned better with the fossil record based dating (7 Mya for MRCA for genus *Pungitius* spp; Rawlinson and Bell, 1982) and with phylogenies based on direct and indirect fossil dating (e.g. Guo et al. 2019). While this provides further confidence to believe that divergence time estimates using our *de novo* mutation rate estimate are closer to the truth than the substitution based estimates, one should keep in mind that mutation rates may evolve over time and/or vary among different lineages (e.g. Pozi and Penna 2022). This variation would naturally influence estimated divergence times. In this perspective, further studies should seek to obtain mutation rate estimates from other members of the family Gasterosteidae.

While leveraging empirically estimated *µ* in divergence time estimation has its advantages (Tiley et al. 2020), one has to remember that the estimated divergences need to be scaled to absolute time units using generation time. Hence, any errors or biases in applied generation time will progate with the divergence time estimates. Since there is considerable variation in life span (three to seven years) and likely also generation time (defined as average age of breeding parents in the population; Hill 1979) among different nine-spined stickleback populations (De Faveri et al. 2014), this is clearly a point of potential concern. However, most of the variation in the generation length occurs between marine (younger) and freshwater (older) populations. Since our estimates of *µ* were derived from marine fish using two-year generation time, we feel confident that generation time is not of concern in this study, or at least when it comes to marine populations. However, since some of the populations and species we analysed originate from freshwater locations, the possibility of some bias cannot be definitely ruled out. Nevertheless, the magnitude of this problem is unlikely to be anywhere close to the difference we observed when estimating divergence times with the synonymous substitution rate as the prior (fig. 4). Moreover, it is also possible that mutation rates in freshwater populations are higher than those in marine populations – if so, this would counteract this problem.

Finally, in spite of the fact that DNM rates are known to be higher in mitochondrial than in the autosomal genome (Nabholz et al. 2008; Xu et al. 2012; Lawless, et al. 2020), we did not detect any mitochondrial mutations in our data. The reason for this is likely to be trivial: Assuming a mutation rate of 1.67 × 10^−8^/bp/yr (an average value of examples in Burridge et al. 2008) and given the size of mitogenome is quite small, with only 16,720 bp for *P. pungitius* (Guo et al. 2016), one would need to survey at least 895 trios to find one mutation in mtDNA. Hence, estimation of mitochondrial mutation rate would require an entirely different sequencing strategy to the one employed in the present study.

In conclusion, the results provide the first and accurate estimate of *µ* for a popular stickleback model system in evolutionary biology. They further show that application of this estimate on divergence time calibration among different stickleback clades pushes back the earlier estimates of divergence times among different lineages, highlighting its utility in phylogenetic and demographic inference. Compared to mutation rate estimates in other vertebrates, the stickleback estimate falls into the middle range, but ranks as the highest reported for teleost fishes depauperate of *µ* estimates. As the estimates in this study came from outbred marine populations, future estimates of *µ* from isolated freshwater populations, as well as from closely related species could provide insights on factors contributing to evolution of mutation rates.

## Data accessibility statement

The raw whole-genome sequencing data and the VCF files will be provided in the European Nucleotide Archive (ENA; https://www.ebi.ac.uk/ena) under accession code PRJEB60682 at the time of acceptance. The nexus alignment files applied in phylogenetic analyses were compiled from Guo et al. (2019) and the .xml control files are available as supplementary files. The annotation file of the reference genome (Version 6) was obtained from Varadharajan *el al*. (2019; https://doi.org/10.6084/m9.figshare.10565507.v1) and the liftover file to Version 7 was published by Rastas (2020; https://sourceforge.net/p/lep-anchor/code/ci/master/tree/liftover.awk). The codes used in the analyses are available on github at the time of acceptance (https://github.com/zcharlene/dnmrate9spinedmarine).

## Acknowledgements

Our research was supported by a grant from the Academy of Finland (#218343 to JM), a grant from the Helsinki Institute for Life Sciences (HiLife; to JM) and a grant from the NSFC/RGC Joint Research Scheme sponsored by the Research Grants Council of the Hong Kong Special Administrative Region, China and the National Natural Science Foundation of China (Project No. N_HKU763/21). C.Z., K.R. and A.F.S. were supported by Faculty of Science (HKU) funding to J.M. We thank Xueyun Feng, Jilong Ma, Dandan Wang and Mikko J Kivikoski for their helpful advice in data analyses. We acknowledge CSC – IT Center for Science, Finland – for access to computational resources and user support.

## Ethic statement

The fish breeding was conducted under a permit from the Animal Experiment Board in Finland (permit reference ESAVI/4979/2018). The parental generation fish from marine sites were collected under national fishing licences.

## Author contributions

Conceived and designed the study: JM, CZ Analysed the data: CZ, KR, MHS, AS Contributed materials/analysis tools: AF, JM Wrote the paper: CZ, JM, KR, AS, MHS

