## supplementary for "*De novo* mutation rates in sticklebacks"

Supplementary materials

**Supplementary Fig. 1. Detected DNMs mapped on *P. pungitius* chromosomes.** Vertical lines depict DNMs and they are coloured according to their genomic location. Included are autosomal linkage groups and pseudoautosomal part (non-grey region) of the linkage group 12 (sex chromosomes).

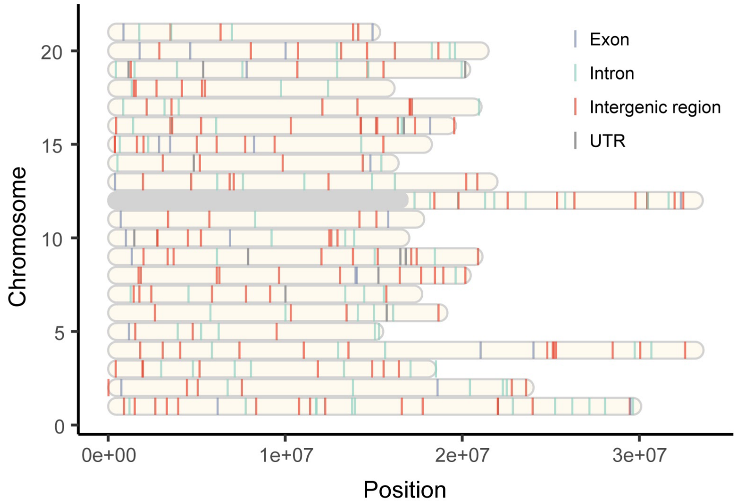

**Supplementary Fig. 2. Power of individual filters across different families.** Number of *de novo* mutation candidates left after being filtered by each individual filter. TVA = Tvärminne family (n = 5), POR = Pori family (n = 4).

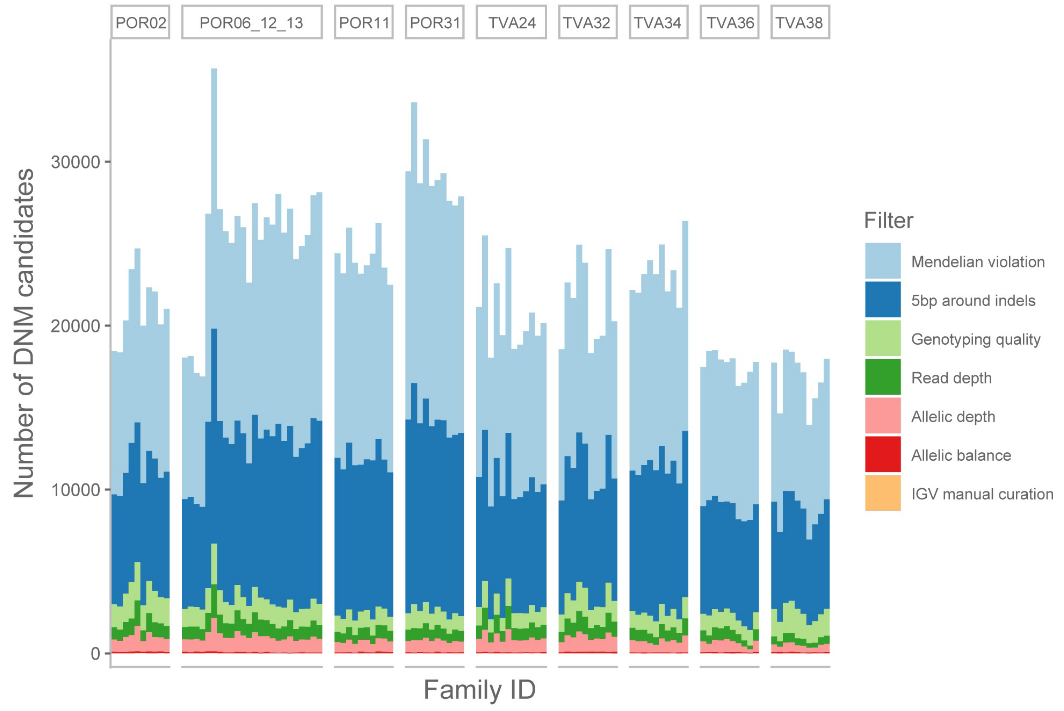


**Supplementary Fig. 3. Comparison of DNM and transmission rates.** There is no statistically significant difference in DNM rates between a) generations, b) sexes, or c) pedigree types. d) Transmission rates of the DNMs from $F_1$ to $F_2$ generation do not differ between inbred and outbred pedigrees.

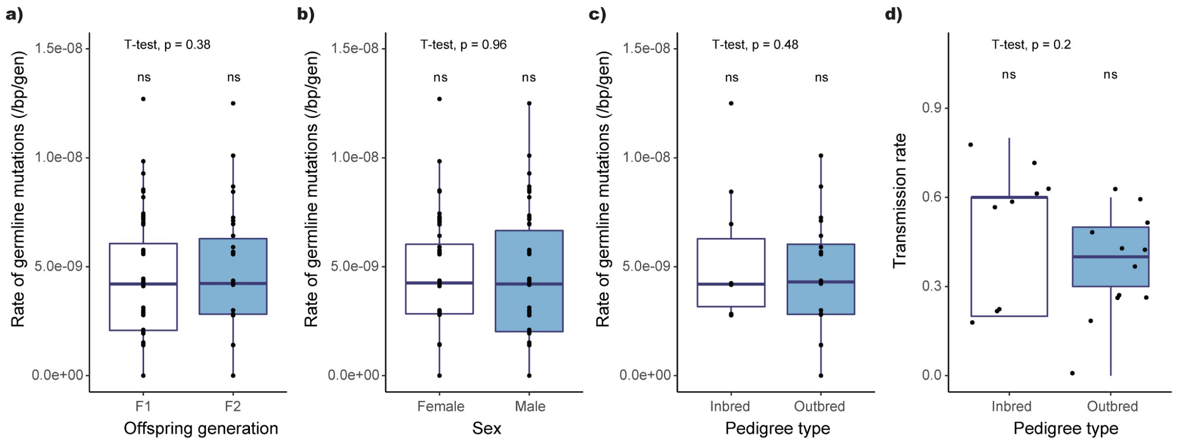

**Supplementary Fig. 4. Detailed DNM- (above) and substitution- (below) rate based phylogenies .** Node A: Divergence time (Mya) between *P. pungitius* and *G. aculeatus.* Node B: Divergence time among different *P. pungitius* lineages. Node C: Divergence time between western and eastern European *P. pungitius* lineages. The 95% HPDs of node ages are indicated by the blue bars. Nodes without white circles are supported by higher than 0.95 posterior values.

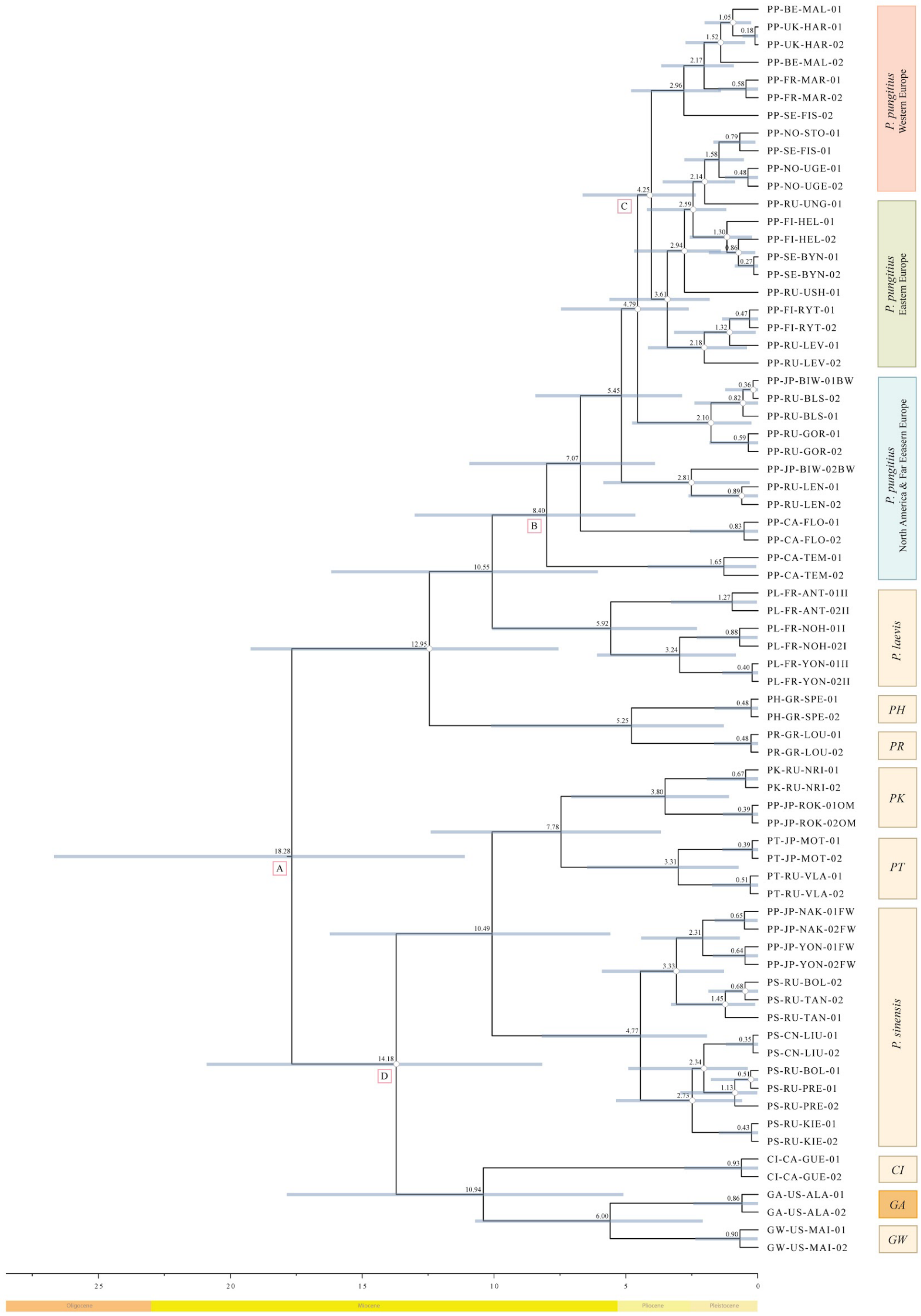

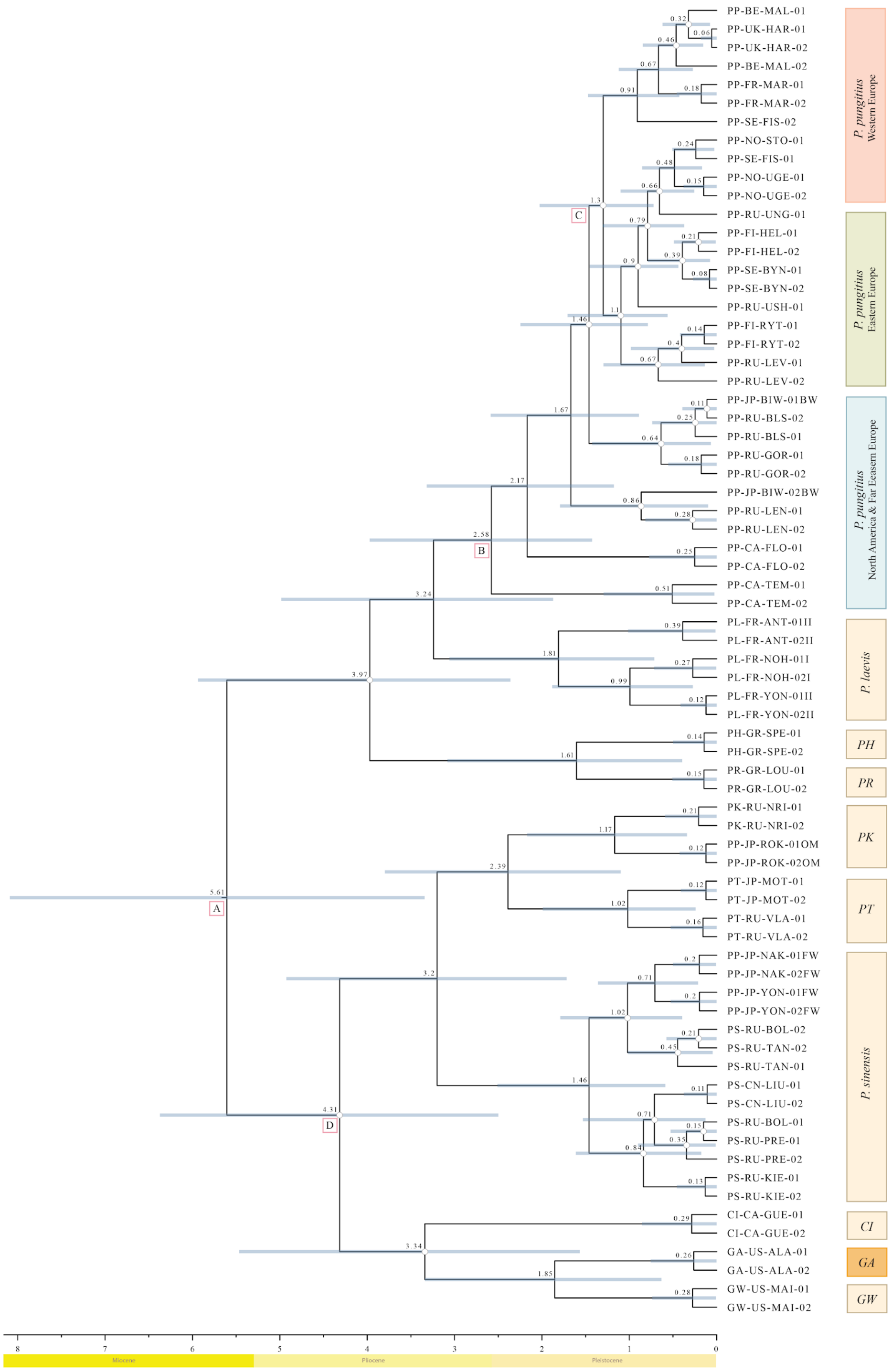
PP-BE-MAL-01
PP-UK-HAR-01
PP-UK-HAR-02
PP-BE-MAL-02
PP-FR-MAR-01
PP-FR-MAR-02
PP-SE-FIS-02
PP-NO-STO-01
PP-SE-FIS-01
PP-NO-UGE-01
PP-NO-UGE-02
PP-RU-UNG-01
PP-FI-HEL-01
PP-FI-HEL-02
PP-SE-BYN-01
PP-SE-BYN-02
PP-RU-USH-01
PP-FI-RYT-01
PP-FI-RYT-02
PP-RU-LEV-01
PP-RU-LEV-02
PP-JP-BIW-01BW
PP-RU-BLS-02
PP-RU-BLS-01
PP-RU-GOR-01
PP-RU-GOR-02
PP-JP-BIW-02BW
PP-RU-LEN-01
PP-RU-LEN-02
PP-CA-FLO-01
PP-CA-FLO-02
PP-CA-TEM-01
PP-CA-TEM-02
PL-FR-ANT-01II
PL-FR-ANT-02II
PL-FR-NOH-01I
PL-FR-NOH-02I
PL-FR-YON-01II
PL-FR-YON-02II
PH-GR-SPE-01
PH-GR-SPE-02
PR-GR-LOU-01
PR-GR-LOU-02
PK-RU-NRI-01
PK-RU-NRI-02
PP-JP-ROK-01OM
PP-JP-ROK-02OM
PT-JP-MOT-01
PT-JP-MOT-02
PT-RU-VLA-01
PT-RU-VLA-02
PP-JP-NAK-01FW
PP-JP-NAK-02FW
PP-JP-YON-01FW
PP-JP-YON-02FW
PS-RU-BOL-02
PS-RU-TAN-02
PS-RU-TAN-01
PS-CN-LIU-01
PS-CN-LIU-02
PS-RU-BOL-01
PS-RU-PRE-01
PS-RU-PRE-02
PS-RU-KIE-01
PS-RU-KIE-02
CI-CA-GUE-01
CI-CA-GUE-02
GA-US-ALA-01
GA-US-ALA-02
GW-US-MAI-01
GW-US-MAI-02
P. pungitius Western Europe
P. pungitius Eastern Europe
P. pungitius North America & Far Eeasern Europe
P. laevis
PH
PR
PK
PT
P. sinensis
CI
GA
GW
A
B
C
D
5.61
4.31
3.97
3.24
3.34
3.2
2.58
2.39
2.17
1.85
1.81
1.67
1.61
1.46
1.3
1.17
1.1
1.02
8
7
6
5
4
3
2
1
0
Miocene
Pliocene
Pleistocene

**Supplementary Table 1. DNM rates estimates from other pedigree-based studies.**

| Species | DNM rates (E-08/generation/bp) | Number of trios | References |
|---|---|---|---|
| Human (*Homo sapiens*) | 1.17 | 1 (CEU) | Conrad et al. 2011 |
| | 0.97 | 1 (YRI) | |
| | 1.2 | 78 | Kong et al. 2012 |
| | 1.27 | 10 | Besenbacher et al. 2015 |
| | 1.28 | 13 | Rahbari et al. 2016 |
| | 1.05 | 719 | Wong et al. 2016 |
| | 1.29 | 1550 | Jónsson et al. 2017 |
| | 1.28 | 150 | Maretty et al. 2017 |
| | 1.7 | 516 | Turner et al. 2017 |
| | 1.1 | 593 | Sasani et al. 2019 |
| | 1.22 | 1449 | Kessler et al. 2020 |
| Chimpanzee (*Pan troglodytes*) | 1.2 | 6 | Venn et al. 2014 |
| | 1.48 | 1 | Tatsumoto et al. 2017 |
| | 1.26 | 7 | Besenbacher et al. 2019 |
| Gorilla (*Gorilla gorilla*) | 1.13 | 2 | |

| | | | |
|---|---|---|---|
| Orangutan (*Pongo abelii*) | 1.66 | 1 | |
| Baboon (*Papio anubis*) | 0.57 | 12 | Wu et al. 2020 |
| Rhesus macaque (*Macaca mulatta*) | 0.58 | 14 | Wang et al. 2020 |
| | 0.77 | 19 | Bergeron et al. 2021 |
| Green monkey (*Chlorocebus sabaeus*) | 0.94 | 3 | Pfeifer, 2017 |
| Owl monkey (*Aotus nancymaae*) | 0.81 | 14 | Thomas et al. 2018 |
| Marmoset (*Callithrix jacchus*) | 0.43 | 1 | Yang et al. 2021 |
| Gray mouse lemur (*Microcebus murinus*) | 1.52 | 2 | Campbell et al. 2021 |
| Mouse (*Mus musculus*) | 0.57 | 8 | Milholland et al. 2017 |
| | 0.39 | 15 | Lindsay et al. 2019 |
| Cattle (*Bos taurus*) | 1.17 | 5 | Harland et al. 2017 |
| Wolf (*Canis lupus*) | 0.45 | 4 | Koch et al. 2019 |
| Boar (*Sus scrofa*) | 0.36 | 5 | Zhang et al. 2022 |
| Domestic cat (*Felis catus*) | 0.86 | 11 | Wang et al. 2022 |
| Platypus (*Ornithorhynchus anatinus*) | 0.7 | 2 | Martin et al. 2018 |
| Collared flycatcher (*Ficedula albicollis*) | 0.46 | 7 | Smeds et al. 2016 |
| Herring (*Clupea harengus*) | 0.2 | 12 | Feng et al. 2017 |

| | | | |
|---|---|---|---|
| Cichlids (*Astatotilapia calliptera*, *Aulonocara stuartgranti*, and *Lethrinops lethrinus*) | 0.35 | 9 | Malinsky et al. 2018 |
| Fruit fly (*Drosophila melanogaster*) | 0.28 | 12 | Keightley et al. 2014 |
| Butterfly (*Heliconius melpomene*) | 0.29 | 30 | Keightley et al. 2015 |
